# Exploring American Elderberry Compounds for Antioxidant, Antiviral, and Antibacterial Properties Through High-Throughput Screening Assays Combined with Untargeted Metabolomics

**DOI:** 10.1101/2024.09.13.611920

**Authors:** Amanda Dwikarina, Mohamed Bayati, Novianus Efrat, Anuradha Roy, Zhentian Lei, Khanh-Van Ho, Lloyd Sumner, Michael Greenlief, Andrew L.Thomas, Wendy Applequist, Andrew Townesmith, Chung-Ho Lin

## Abstract

American elderberry (*Sambucus nigra* subsp. *canadensis*) is a rapidly emerging new perennial crop for Missouri, recognized for its high level of bioactive compounds with significant health benefits, including antibacterial, antiviral, and antioxidant properties. A high-throughput screening assay combined with untargeted metabolomics analysis was utilized on American elderberry juice from 21 genotypes to explore and characterize these bioactive compounds. Our metabolomics study has identified 32 putative bioactive compounds in the American Elderberry juices. An array of high-throughput screening bioassays was conducted to evaluate 1) total antioxidant capacity, 2) activation of antioxidant response elements (ARE), 3) antiviral activity, and 4) antibacterial activity of the putatively identified compounds. Our results revealed that 14 of the 32 American elderberry compounds exhibited strong antioxidant activity. Four compounds (isorhamnetin 3-O-glucoside, kaempferol, quercetin, and naringenin) activated ARE activity and were found to be non-cytotoxic to cells. Notably, six of the 32 compounds demonstrated significant antiviral activity in an *in vitro* TZM-bl assay against two strains of HIV-1 virus, CXCR4-dependent NL4-3 virus and CCR5-dependent BaL virus. Luteolin showed the most potent anti-HIV activity against the NL4-3 virus (IC_50_ = 1.49 µM), followed by isorhamnetin (IC_50_ = 1.67 µM). The most potent anti-HIV compound against the BaL virus was myricetin (IC_50_ = 1.14 µM), followed by luteolin (IC_50_ = 4.38 µM). Additionally, six compounds were found to have antibacterial activity against gram-positive bacteria *S. aureus*, with cyanidin 3-O-rutinoside having the most potent antibacterial activity in vitro (IC_50_ = 2.9 µM), followed by cyanidin 3-O-glucoside (IC_50_ = 3.7 µM). These findings support and validate the potential health benefits of compounds found in American elderberry juices and highlight their potential for use in dietary supplements as well as innovative applications in health and medicine.

## 1. Introduction

Elderberries are perennial fruit-producing shrubs or small trees belonging to the family Viburnaceae and genus *Sambucus*^1^. *Sambucus nigra* subsp. *nigra* (European elderberry) and *Sambucus nigra* subsp. *canadensis* (American elderberry) are two taxa of elderberry that are increasingly being commercially produced. American elderberry is native to much of eastern and midwestern North America and has been used for both food and medicine by indigenous communities for millennia^2–4^. It is emerging as an important specialty crop in the midwestern USA, mostly cultivated for processing markets^5^. Both fruits and flowers have become increasingly popular for use in a variety of foods, wines, and health supplements^6,7^.

Elderberry fruits and flowers contain abundant health-promoting bioactive compounds, including phenolics, flavonoids, and antioxidant phytochemicals^8^. Several studies analyzed the bioactive composition of American and European elderberries, identifying different polyphenolic compounds in the fruits, flowers, leaves, and stems^9–12^. Elderberry’s major phenolic and flavonoid compounds include gallic acid, neochlorogenic acid, rutin, and quercetin^9–12^. In addition, elderberry fruits contain large amounts of anthocyanins, such as cyanidin-3-glucoside and cyanidin-3-sambubioside, which exhibit antioxidant activity ^8,13^.

Basic research has demonstrated the potential health benefits of elderberry, showcasing a wide range of biological functions, including antioxidant^10,11,14,15^, anti-inflammatory^13,16,17^, anticancer^18,19^, antiviral^8,20^, antibacterial^15,21^, and anti-diabetic^22^ attributes. However, most research has focused on European elderberry, with only limited studies on the health-promoting activities of American elderberry. Modern technology, such as high-throughput screening bioassays and metabolomic analyses, can be used to untangle and elucidate the potential health-benefitting metabolites of American elderberry at much higher levels than have been previously possible.

High-throughput screening (HTS) is a subset of the drug discovery process that incorporates the multidisciplinary fields of analytical chemistry, biology, biochemistry, and computational technologies. It plays an important role, especially when novel biochemical targets are involved^23^. The HTS allows for a standardized, rapid, and cost-effective evaluation of small molecules and novel chemical biological functions. In recent years, HTS labs have used a combination of biochemical and cell-based assays. Both are necessary for discovering and characterizing newly identified compounds in the drug discovery process^23,24^.

This project combined high-throughput screening assays and metabolomics analyses to establish a standardized strategy for investigating the biological functions of American elderberry, including its antioxidant, antibacterial, and antiviral properties. With advancements in computational power and database capacity, our objective was to explore the bioactive activities of American elderberry and facilitate the development of new economic opportunities.

## 2. Materials and Method

### 2.1 Plant material

Elderberry fruits from 21 American elderberry (*Sambucus nigra* subsp. *canadensis*) accessions and cultivars (Table) were harvested from plantings at the University of Missouri’s Southwest Research, Extension, and Education Center near Mt. Vernon, Missouri, U.S.A. The samples represent wild American elderberry accessions from across the midwestern and eastern USA, including collections amassed and studied in Mudge et al. (2016)^25^, as well as established cultivars (Byers et al. 2010; Byers and Thomas 2011). For simplicity, all elderberry accessions and cultivars described in this study are henceforth noted as “genotypes”. Elderberry fruit was harvested at peak ripeness and immediately frozen. Later, the fruit was de-stemmed, thawed at ambient temperature, and the juice pressed by hand, filtered through a kitchen sieve, aliquoted into 50 mL polypropylene tubes, and re-frozen (−20 °C) until further processing.

**Table 1.** American elderberry accessions and cultivars (genotypes) evaluated in this study.

| Accession/Cultivar | Collector<sup>a</sup> | State (USA) | Accession/Cultivar Citations |
| --- | --- | --- | --- |
| 1069 | Meyer | Pennsylvania | Mudge et al. 2016<sup>25</sup> |
| 1076 | Meyer | New York | Mudge et al. 2016<sup>25</sup> |
| 1191 | Meyer | Arkansas | Mudge et al. 2016<sup>25</sup> |
| 1196 | Meyer | Alabama | Mudge et al. 2016<sup>25</sup> |
| 1199 | Meyer | Georgia | Mudge et al. 2016<sup>25</sup> |
| 1887A | Townesmith | West Virginia | Mudge et al. 2016<sup>25</sup> |
| 1889A | Townesmith | West Virginia | Mudge et al. 2016<sup>25</sup> |
| 1892 | Townesmith | New York | Mudge et al. 2016<sup>25</sup> |
| 1896 | Townesmith | Connecticut | Mudge et al. 2016<sup>25</sup> |
| 1911 | Townesmith | Pennsylvania | Mudge et al. 2016<sup>25</sup> |
| 1913 | Townesmith | Pennsylvania | Mudge et al. 2016<sup>25</sup> |
| 2073 | Townesmith | Mississippi | Mudge et al. 2016<sup>25</sup> |
| 2079 | Townesmith | South Carolina | Mudge et al. 2016<sup>25</sup> |
| 2083 | Townesmith | South Carolina | Mudge et al. 2016<sup>25</sup> |
| 2084 | Townesmith | South Carolina | Mudge et al. 2016<sup>25</sup> |
| 2085 | Townesmith | South Carolina | Mudge et al. 2016<sup>25</sup> |
| 2089 | Townesmith | North Carolina | Mudge et al. 2016<sup>25</sup> |
| 2095 | Townesmith | West Virginia | Mudge et al. 2016<sup>25</sup> |
| Bob Gordon (BG) | Thomas | Missouri | Byers and Thomas 2011<sup>26</sup> |
| Ozark | Byers | Arkansas | Thomas et al. 2015<sup>6</sup> |
| Wyldeewood (WW) | Millican | Oklahoma | Byers et al. 2010<sup>27</sup> |

<sup>a</sup>Accession/cultivar collectors are Karen M. Meyer and Andrew K. Townesmith (Missouri Botanical Garden, St. Louis, MO), Andrew L. Thomas and Patrick L. Byers (University of Missouri, Columbia, MO), and Margaret Millican (Wyldeewood Cellars, Wichita, KS).

### 2.2 Sample preparation

All American elderberry juice samples were thawed, then centrifuged at 8000 rpm for 20 minutes. One mL of supernatant was mixed with four mL of HPLC-grade methanol (Fisher Scientific, PA, USA). The mixture was then sonicated for 60 minutes using a sonicator (Fischer Scientific FS-60). The mixture was then filtered through a 0.2 μm PTFE Acrodisc syringe filter (Waters, USA) to remove solid components and transferred to an HPLC autosampler vial (Waters, USA) for further analyses.

### 2.3 Untargeted metabolomics analyses

The untargeted metabolomics profiling was performed on a Bruker maXis impact II quadrupole-time-of-flight (Q-ToF) mass spectrometer (MS) coupled with Waters Acquity Ultra Performance Liquid Chromatography (UPLC). High-resolution mass spectrometry was performed in negative and positive electrospray ionization mode as a preliminary experiment to identify the chemical compounds in the plant extract. Separation was achieved on a Waters C18 column (2.1 × 150 mm, BEH C18 column with 1.7 µm particles) using a linear gradient of 95%:5% to 30%:70% eluent A:B (A: 0.1% formic acid in water, B: acetonitrile) over 30 min. Between 30 – 33 min, the linear gradient was increased from 70% to 95% B and maintained at 95% B for 3 min. The percentage of B was maintained at 5% from 36 – 40 min. The flow rate was 0.56 mL/min, and the column temperature was 60°C. Mass spectrometry was performed in the positive and negative electrospray ionization mode with the nebulization gas pressure at 43.5 psi, dry gas of 12 L/min, dry temperature of 250°C, and a capillary voltage of 4000 V. Auto MS/MS tandem mass spectral data were collected using the following parameters: MS full scan: 100 to 1500 m/z; the number of precursors for MS/MS: 3; threshold: 10 counts; active exclusion: 3 spectra, released after 0.15 min; collision energy: dependent on mass, 10eV at 50 Da, 20 eV at 200 Da, 30 eV at 500 Da, 40 eV at 1000 Da, and 50 ev at 1500 Da. The MS and MS/MS data were auto-calibrated using sodium formate that was introduced at the end of the gradient and after data acquisition.

### 2.4 Metabolomics data analyses

Bioactive compounds were identified in elderberry juice using the web-based platform XCMS Online, which is equipped with the METLIN database and is freely available at https://xcmsonline.scripps.edu. The METLIN library contains over one million molecules to facilitate metabolite identification. UHPLC-MS chromatograms were aligned using the R package XCMS. CentWave settings were used for feature detection (maximal tolerated m/z deviation = 10 ppm, minimum peak width = 5 s, and maximum peak width = 20 s). Obiwarp settings were used for retention time correction (profStep = 1). The chromatogram alignment setting for peak alignment was as follows: bw = 5, minfrac = 0.5, mzwid 0.015. The statistical tests are carried out systematically following feature detection and profile alignment using the Welch t-test (p-value <0.05).

Metabolite UHPLC-MS/MS mass spectral data were analyzed using MS-Dial Software (http://prime.psc.riken.jp/) to further aid compound identification. Data files were converted to ABF format using AnalysisBaseFile Converter (https://www.reifycs.com/AbfConverter/) before import to MS Dial software. Peak picking, alignment, deconvolution, and MS/MS database search were performed with the following parameter: MS1 tolerance 0.01 Da, MS2 tolerance 0.05 Da, identification score cut off 70%, minimum peak height 3000, mass slice width 0.1 Da, retention time tolerance 0.05 min, MS/MS databases ESI(+) and (-)-MS/MS from authentic standards. The relative abundance of putative metabolites identified were further statistically analyzed using the web-based platform MetaboAnalyst ver. 6.0 (https://www.metaboanalyst.ca/).

### 2.5 High-throughput screening assay (HTS)

Our metabolomics analysis putatively identified 32 phenolic compounds in American elderberry juices. These compounds were further evaluated for their antioxidant, anti-HIV, and antibacterial activities using a high-throughput screening assay (HTS) at the Kansas High Throughput Screening Laboratory at the University of Kansas, Lawrence, KS (https://hts.ku.edu/).

#### 2.5.1 Total antioxidant capacity

The total antioxidant activity of American elderberry compounds was evaluated using an electron transfer-based Cupric Reducing Antioxidant Capacity (CUPRAC) method^28^. Briefly, the compounds were solvated in 100% DMSO and transferred acoustically to prepare assay plates using ECHO 655 (Beckman Inc.). The compounds were tested in a 7-concentration dose response in 384 well microplates. The compounds were mixed with 50 µl of freshly prepared CUPRAC reagent (Cu(II)-neocuproine. After mixing at 650 rpm for 2 mins, the plates were sealed and incubated for up to 3 h at 25°C. The absorbance of the samples of Cu(I)-neocuproine (Nc) chelate formed from redox reaction between antioxidants and CUPRAC reagent was read at 1.5 h and 3 h at 450 nm using a microplate reader (BioTek Neo, Agilent). A Trolox standard curve was included on each assay plate, and the total antioxidant capacity of the compounds was interpolated and expressed as Trolox equivalent (mM) from the Trolox standard curve of the Trolox control.

For total antioxidant capacity analysis, linear regression analysis was performed using GraphPad Prism 8 (San Diego, CA, USA) to identify the linear regression equation for each compound. The coefficient of the compound equation was compared with the coefficient of the Trolox control to determine each compound’s relative total antioxidant capacity. The fold increase over Trolox was calculated by dividing the compound models’ coefficient by the Trolox control coefficient. The compounds that exhibited a fold increase over Trolox greater than 5 were considered to possess significant total antioxidant capacity.

#### 2.5.2 Antioxidant Response Element (ARE) activation

American elderberry compounds were evaluated for Nuclear factor erythroid 2-related factor 2 (Nrf2) activation using ARE-Luciferase reporter-based assay in the HepG2 cell line^29^. The method for ARE activation assay is as follows: a HepG2 hepatic cell line stably expressing firefly luciferase (Luc) reporter under the control of ARE was obtained from BPS Bioscience. Briefly, the HepG2-ARE-Luc cells (10,000 cells/well) were seeded in 384-well plates in 50 µL complete media per well using a Multidrop Combi dispenser (Thermo Fisher Scientific, Waltham, MA, USA). Compounds (7-concentration dose-response) were transferred to the plates and incubated at 37 °C in a 5% CO_2_ humidified incubator for 18 h. Each assay plate also contained known positive controls [tert-butylhydroquinone (TBHQ) and DL-Sulphorane] as well as DMSO (as a vehicle negative control). The reporter activity was measured by the addition of 25 µL Steady-Glo^®^ luciferase assay reagent (Promega Inc.) for 30 min. The luminescence intensities of the 384-well plates were read on a BioTek Neo microplate reader (Agilent Inc.). The increase in luciferase activity correlates with compound-induced activation of Nrf2. The fold increase in luciferase was normalized to the positive and negative controls on each assay plate.

The ARE fold induction of the compounds was measured by dividing the luminescence absorbance of the treatment by the specific luminescence absorbance of the control sample and multiplying by 100. The control sample (in the presence of DMSO vehicle and without the compounds) was set at 100%. The compounds with ARE fold induction to 10-fold over the vehicle controls in one or more concentrations were considered to have significant ARE induction activity. The bioactive compounds’ relative cytotoxicity (%) was calculated by dividing the specific luminescence absorbance of the treated sample by the specific luminescence absorbance of the control sample and multiplying by 100. The control sample (in the presence of DMSO vehicle and without the compounds) was set at 100%. Non-linear regression analysis of data was performed to identify the dose-response curve for each compound. The IC_50_ values (half maximal inhibitory concentration) of each compound were determined from the dose-response curve for the A549 and MRC-5 cell lines using GraphPad Prism 8. The compounds that exhibited IC _50_ values < 10 µM in the A549 cell line and had no toxic effects on the control cell line MRC-5 were considered to be potent antiproliferative compounds.

#### 2.5.3 Evaluation of Anti-HIV activity

The anti-HIV activity of 32 putative compounds identified in American elderberry juices was evaluated at the Kansas High Throughput Screening Laboratory at the University of Kansas, Lawrence, KS (https://hts.ku.edu/). The method for anti-HIV activity assay provided by Dr. Anuradha Roy is as follows: all reagents for HIV neutralization assay were obtained through the HIV Reagent Program, Division of Acquired Immunodeficiency Syndrome (AIDS), National Institute of Allergy and Infectious Diseases (NIAID), NIH. TZM-bl reporter cells, derived from Hela cell clone expressing CD4 glycoprotein, CC-chemokine receptor type 5 (CCR5), and CXC-chemokine receptor 4 (CXCR4) as well as an HIV-trans-activator of transcription (HIV-Tat) regulated firefly luciferase & β-galactosidase reporters (Cat. no. 8129), used to optimize the HIV-1 neutralization assay according to reference^30^. Briefly, the TZM-bl cell line was grown at 37°C in humidified 5% CO_2_ in Dulbecco’s Modified Eagle’s Medium with L-glutamine, sodium pyruvate, glucose, and 10% heat-inactivated fetal bovine serum (FBS) and 50 µg gentamicin/ml. Two HIV-1 strains, NL4-3 [ARP-114, X4 tropic] and Bal [ARP-11414, R5 tropic] viruses, were prepared by transfecting 293T/17 cells with DNA constructs containing full-length wild type (WT) sequences, as described previously^31^. The virus was titrated in TZM-bl cells to achieve an average tissue culture infectious dose (TCID) of 150,000 relative light unit (RLU) equivalents for pNL4-3 and TCID of 20,000 for pBAl WT corresponding to 8-10 times the background, respectively.

To identify compounds that interfere with viral replication, 5000 cells/well were seeded in 384 well plates containing compounds. An equal volume of virus (2X stock) was added to the cells in 8 µg/ml diethylaminoethanol (DEAE)-Dextran. After 48h at 37°C, 5% CO_2_, Promega Steady Glo Luciferase detection reagent was added to detect Firefly luminescence. Percent inhibition was normalized to DMSO controls. Four known HIV inhibitors (AZT, JM2987, TAK779 and Maraviroc) were used as positive controls

#### 2.5.4 Antibacterial growth assay

The activity of American elderberry compounds was tested against *Staphylococcus aureus* subsp. *aureus* Rosenbach, ATCC 29213, *Pseudomonas aeruginosa*, PAO1, ATCC 15692, and *Klebsiella pneumoniae* subsp. *pneumoniae*, ATCC 27736. The bacteria from a single colony were grown overnight in Mueller Hinton cation-adjusted broth at 37°C, 250 rpm. The overnight culture was diluted 100-fold, and 40 µl of bacterial suspension was grown in the presence of American elderberry compounds (7 concentration dose-response) for 24h at 37°C. The activity of compounds was also evaluated against *Mycobacterium smegmatis*, which was grown in Middlebrook 7H9 OADC supplemented media at 37°C, 250 rpm, for 24h and 48h. Positive controls included Vancomycin and Amikacin, while DMSO was the negative control on each assay plate. The plates were read at Abs 600 nm at 24h and 48h. After the absorbance reads, Bact-Titer Glo was added to the plates, and an ATP-based luminescence assay was read using BioTek Synergy. Percent inhibition of growth was normalized to positive and negative controls on each assay plate.

## 3. Results

### 3.1 Untargeted metabolomic analysis

Global profiling using UHPLC-QTOF-MS analysis resulted in the putative identification of more than 100 compounds (Supplementary Table 1). Partial least-square discriminant analysis (PLS-DA) showed significant differences in the metabolomics profiles of American elderberry genotypes (Figure 1). PLS-DA analysis with two principal components (PCs) covered 43.5% of the total variability of the data. The first principal component (PC1) explained 20.1% of the data variability, whereas the second principal component (PC2) accounted for 23.4% of the total variability of the data set. The top 25 important compounds putatively identified in American elderberry were identified using the PLS-DA loading plot (Figure 1A). K-means clustering analysis was performed with three random clusters to compare the metabolomic profile of each American elderberry genotype (Figure 1B). The K-means analysis (Supplementary Table 2) revealed that the American elderberry genotype Ozark had a significantly different metabolomic profile than the rest of the American elderberry genotype (Figure 2). The metabolite profile of ‘Ozark’ also differed from other elderberry genotypes studied in Thomas et al. (2015)^6^.

**Figure 1.**
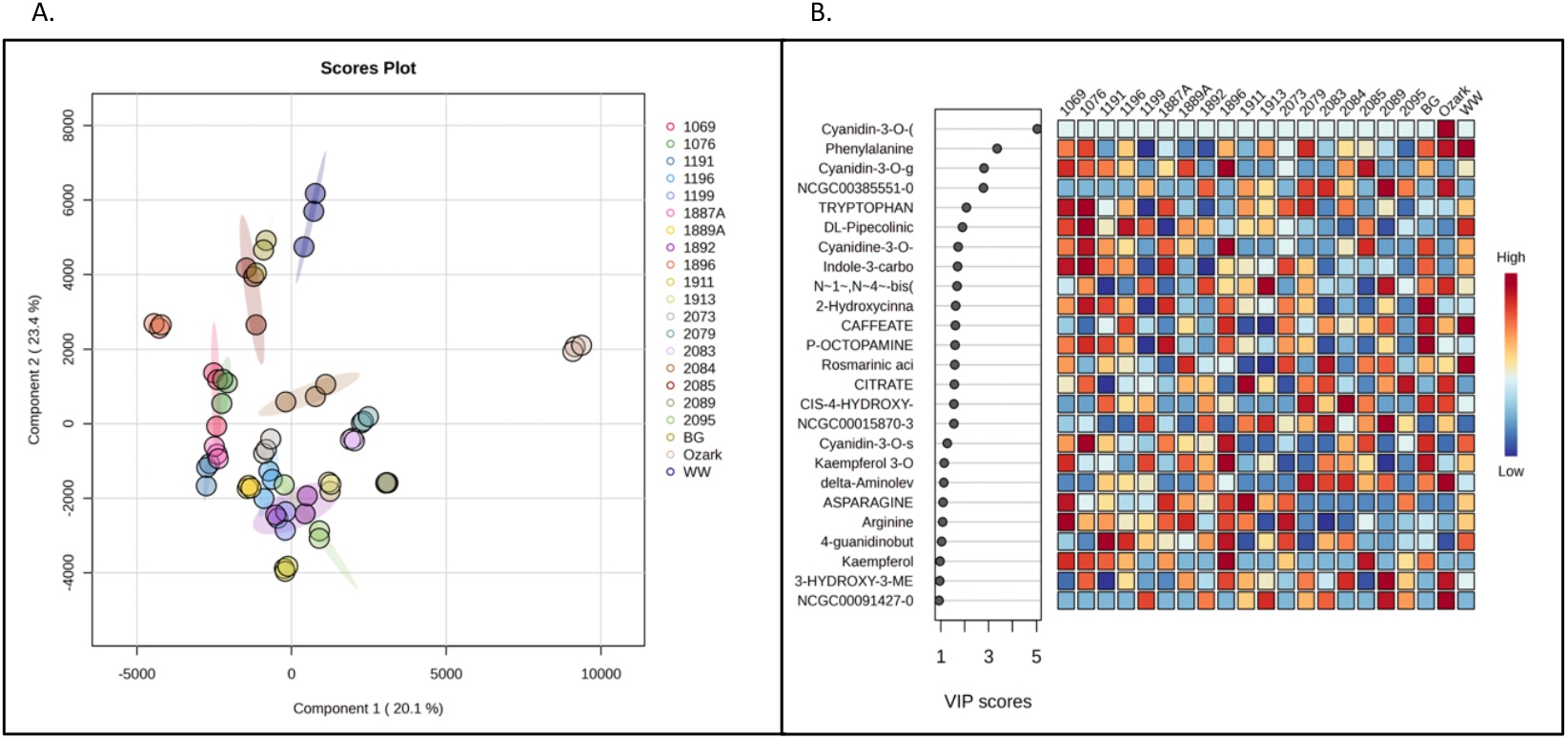
Partial least-squares discriminant analysis (PLS-DA). A. PLS-DA loading plot of putatively identified compounds in juices of 21 American elderberry genotypes revealed differences in the metabolomic profiles. Circles with the same color represent analytical replicates (n=3). The colored ellipses indicate 95% confidence regions of each group. B. Important features identified by PLS-DA. The colored boxes on the right indicate the relative abundance of the corresponding metabolite in each American elderberry genotype under study, with dark red representing high abundance and dark blue representing low abundance.

**Figure 2.**
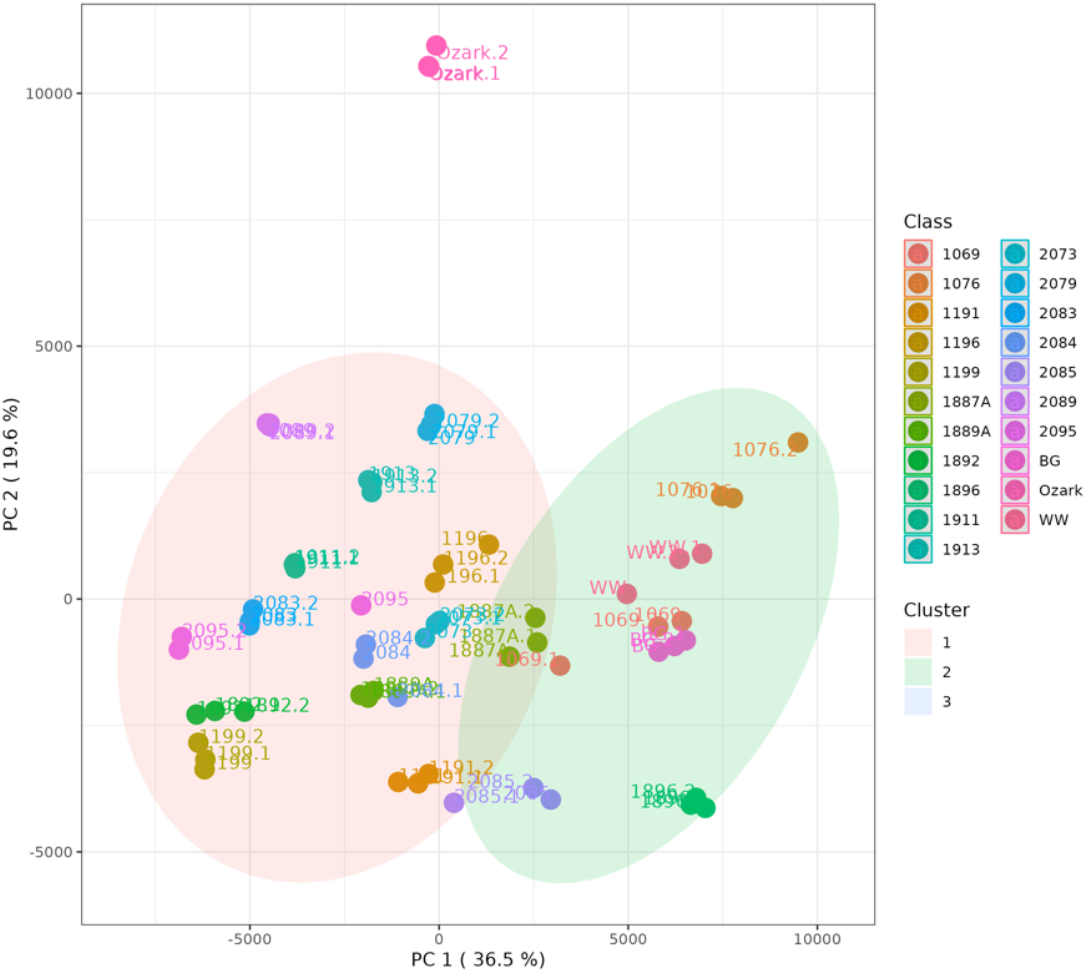
K-mean clustering analysis (cluster = 3) showed a significant difference in the metabolomic profile of the American elderberry genotype. Genotypes with similar metabolomic profiles were clustered together.

The metabolomics analysis resulted in more than 100 putative compounds identified. The heatmap relative intensity of 108 identified compounds in the juices of 21 American elderberry genotypes are presented in Supplementary Figure 1. A comprehensive literature review was conducted to determine the potential biological functions of each compound (Supplementary Table 1.). The 32 chemicals with known bioactive functions were selected for further analysis based on their unique properties for American elderberry (Table 2). A variety of cyanidins, such as cyanidin 3-*O*-sophoroside, cyanidin 3-*O*-sambubioside, cyanidin-3-*O*-rutinoside, and cyanidin 3-*O*-galactoside, were putatively identified in American elderberry juice. A variety of quercetin compounds were also putatively identified in American elderberry juice. These compounds include quercetin, isoquercetin, quercetin-3-rutinoside, quercetin-3-glucoside, and quercetin-3-galactoside. Other than cyanidin and quercetin, other polyphenol compounds, such as isorhamnetin, kaempferol, luteolin, and myricetin, were putatively identified in American elderberry juice. Untargeted metabolomics data showed a variety of specialized metabolites in American elderberry juice whose relative abundance varied across American elderberry genotypes, presented in the heatmap (Figure 3); the darker the color red, the higher the relative abundance of compounds.

**Table 1.**
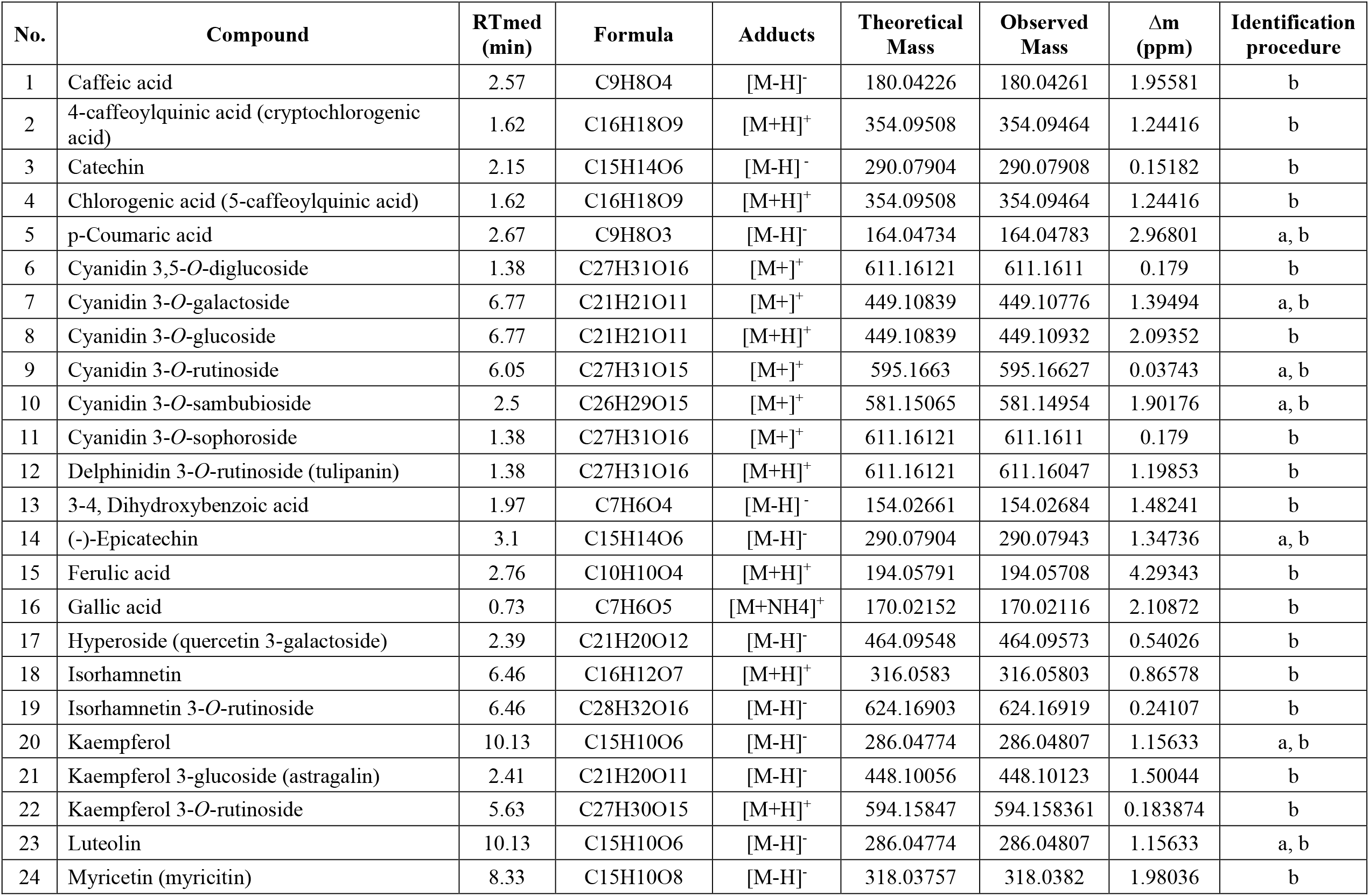
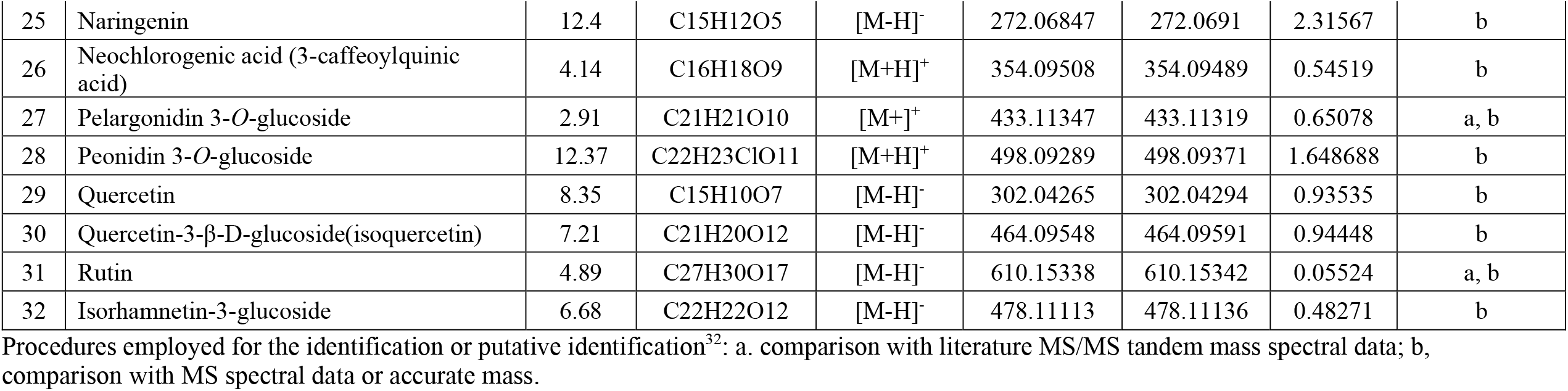
Identification of specialized metabolites in American Elderberry juices through untargeted metabolomics analysis.

**Figure 3.**
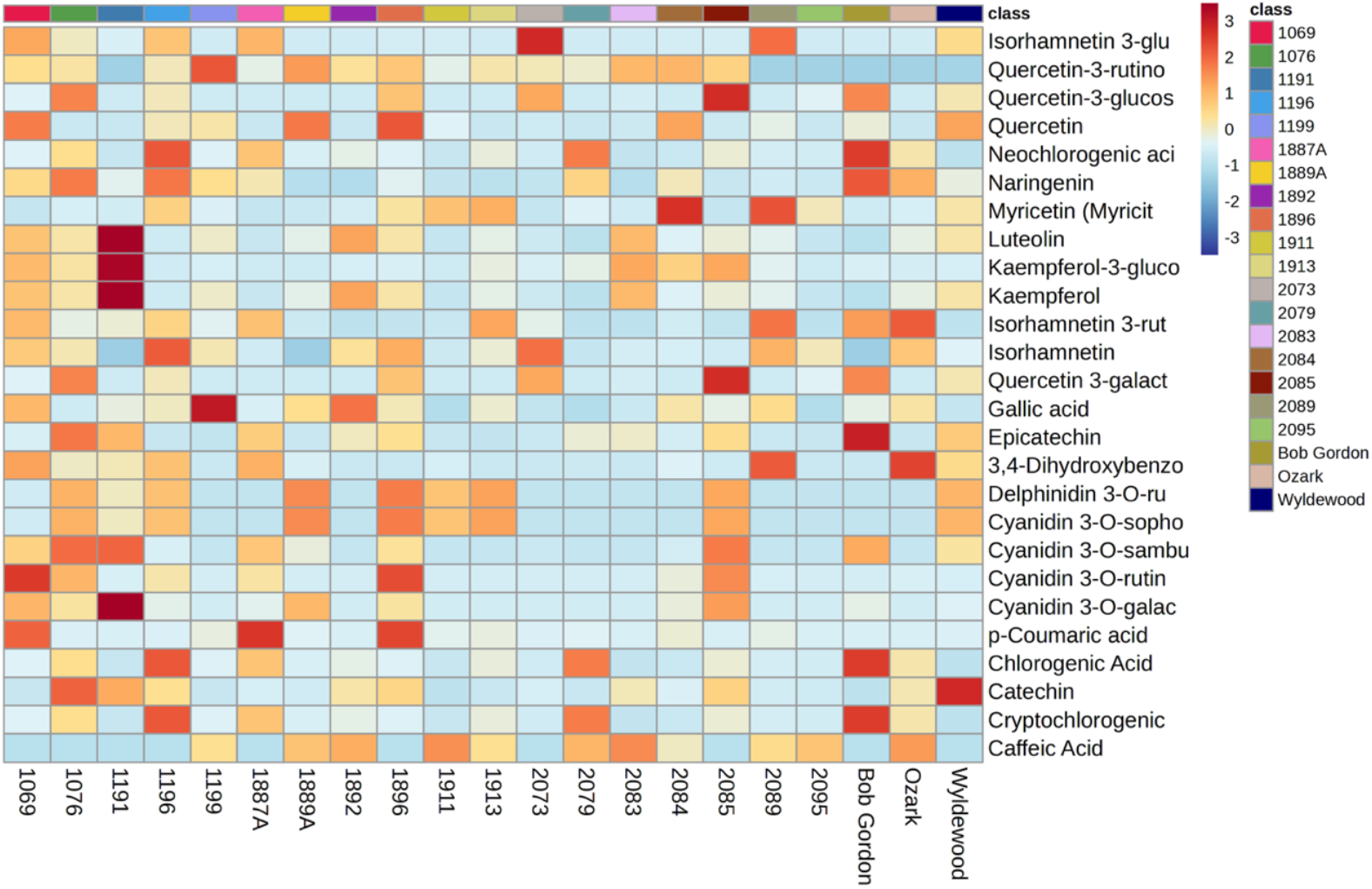
The heatmap shows the relative abundance of 32 selected putatively identified compounds in the juices collected from various American elderberry genotypes. Each color represents a relative abundance of compounds in the juices. The scale ranges from dark red to dark blue, representing the range of high to no abundance of compounds. Heatmap features the top thirty-two metabolite features as identified by t-test analysis (p < 0.05 and intensity ≥10,000). The distance measure is by Euclidean correlation, and clustering is determined using the Ward algorithm.

Compounds quercetin-3-glucoside and quercetin-3-galactoside were detected with the highest relative abundance in genotype 2085. Gallic acid was detected with the highest relative abundance in genotype 1199. The specialized metabolites in Bob Gordon are 4-caffeoylquinic acid, chlorogenic acid, neochlorogenic acid, and epicatechin. Wyldewood contained catechin as its major component. In contrast, the major components of Ozark are isorhamnetin 3-rutinoside and 3,4-dihydroxybenzoic acid. Genotype 1191 contained the highest relative abundance of cyanidin-3-*O*-galactoside, kaempferol, kaempferol 3-glucoside, and luteolin.

Untargeted UHPLC-QTOF-MS-MS data of American elderberry juices were analyzed using MS-DIAL platform. Our results showed 10 putative compounds with matching MS-MS reference spectra (Figure 4). Based on the untargeted UHPLC-QTOF-MS-MS data, putatively identified compounds were catechin, p-coumaric acid, cyanidin-3-*O*-galactoside, cyanidin-3-o-rutinoside, cyanidine-3-*O*-sambubioside, 3,4-dihydroxymandelic acid, epicatechin, isorhamnetin-3-o-rutinoside, kaempferol, luteolin, pelargonidin-3-*O*-glucoside, and quercetin-3-*O*-rutinoside (rutin). Untargeted UHPLC-QTOF-MS-MS data provides additional information for the putatively identified compounds, adding another layer of confirmation.

**Figure 4.**
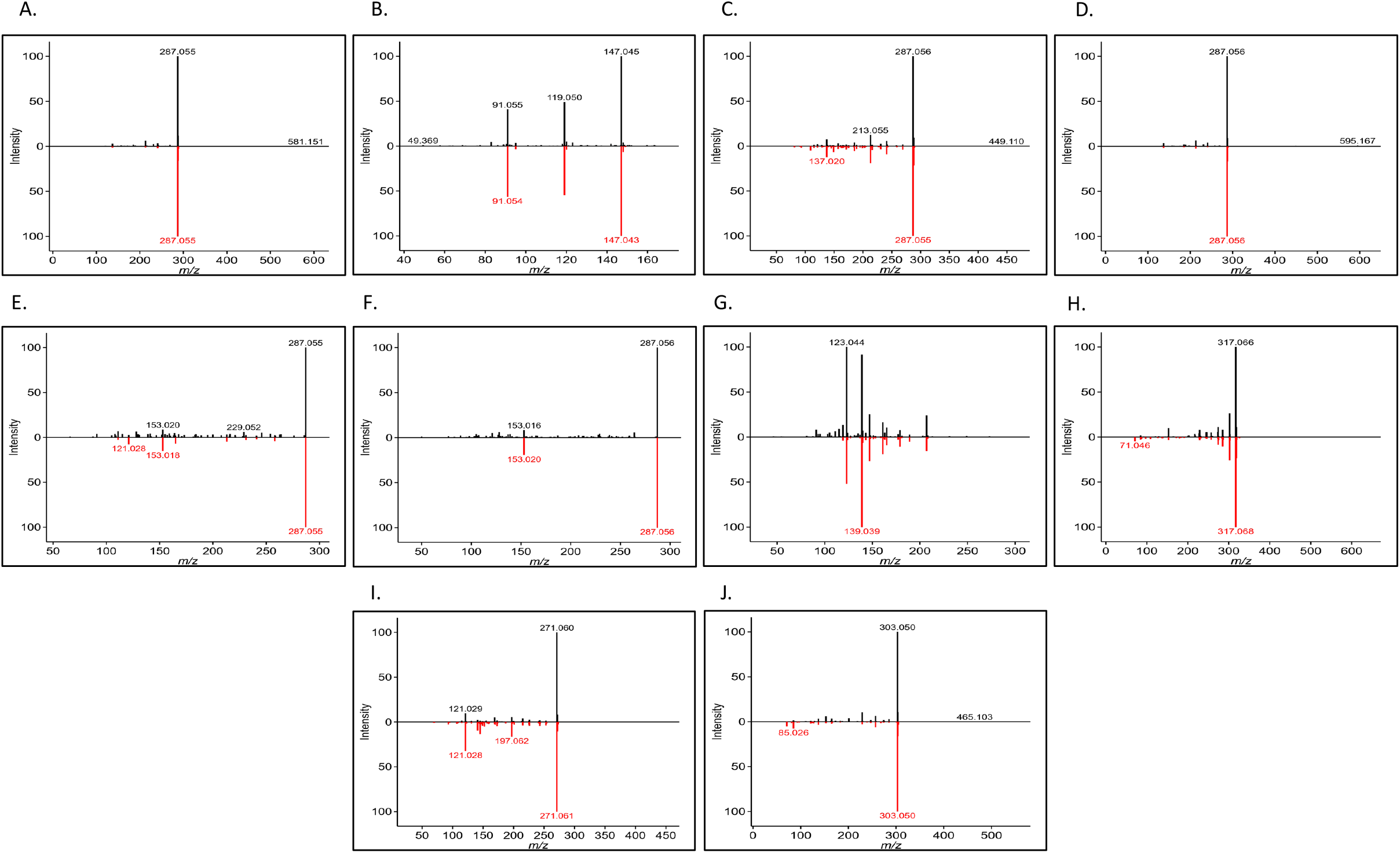
Experiment spectrum (in black) versus reference MS/MS tandem spectrum data (in red). A. Cyanidin-3-*O*-sambubioside, B. p-Coumaric acid, C. Cyanidin-3-*O*-galactoside, D. Cyanidin-3-*O*-rutinoside, E. Kaempferol, F. Luteolin, G. Epicatechin, H. Isorhamnetin-3-*O*-rutinoside, I. Pelargonidin-3-*O*-glucoside, J. Quercetin-3-*O*-rutinoside (Rutin).

### 3.2 Antioxidant compounds in American Elderberry

Putatively identified compounds in American elderberry juice were screened for antioxidant activity at the University of Kansas High Throughput Screening laboratory, Lawrence KS (https://hts.ku.edu/). The total antioxidant capacity (TAC) of American elderberry compounds was performed using CUPRAC methods^28^ with Trolox as control. The total antioxidant capacity was expressed as Trolox equivalent, and compounds that showed TAC values greater than 5 Trolox units at any one concentration tested were considered significant. From 32 compounds tested, 14 compounds showed TAC five-fold greater than Trolox, which are hyperoside (quercetin 3-galactoside), cyanidin 3-*O*-sambubioside, quercetin, cyanidin 3-*O*-galactoside, isorhamnetin, myricetin, delphinidin 3-*O*-rutinoside, cyanidin 3-*O*-sophoroside, cyanidin 3-*O*-rutinoside, cyanidin 3-*O*-glucoside, quercetin-3-β-D-glucoside (hirsutrin), cyanidin 3,5-*O*-diglucoside, neochlorogenic acid (3-caffeoylquinic acid), and gallic acid (Figure 5A).

**Figure 5.**
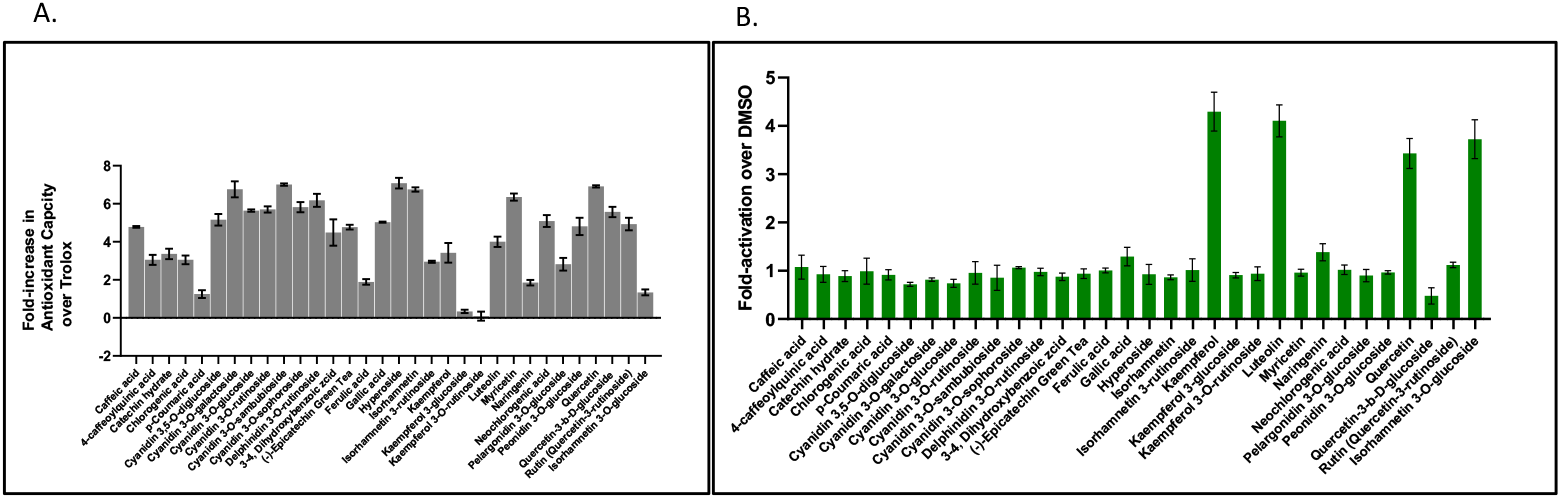
Antioxidant activity result. A. Total antioxidant capacity of compounds in American elderberry juices. Data was expressed as Trolox equivalent (n=2). B. Evaluation of American elderberry compound (at a concentration of 10 µM) on activating antioxidant response element (ARE) in HepG2-Nrf2 reporter cell line (n=2).

Of the 32 compounds putatively identified in American elderberry juices, 12 showed TACs ranging from two to 4.9 times greater than Trolox. These compounds are rutin, peonidin 3-*O*-glucoside, caffeic acid, epicatechin, 3,4-dihydroxybenxoic acid, luteolin, kaempferol, catechin, 4-caffeoylquinic acid (cryptochlorogenic acid), chlorogenic acid, isorhamnetin 3-rutinoside, and pelargonidin 3-*O*-glucoside. Ferulic, acid, naringenin, isorhamnetin 3-*O*-glucoside, p-coumaric acid, kaempferol 3-glucoside, and kaempferol 3-*O*-rutinoside showed TACs on the same range as Trolox.

Putatively identified compounds in American elderberry juices were also evaluated for their ability to activate the Nrf2 transcription factor-ARE signaling pathway in the HepG2 cell line. Four American elderberry compounds activated ARE activity. At a concentration of 10 µM, kaempferol exhibits the highest level of ARE activation, followed by luteolin, isorhamnetin 3-*O*-glucoside, and quercetin, respectively (Figure 5B). All four compounds were found to be non-cytotoxic to the cells (Supplementary Figure 2). Despite their ability to activate the ARE pathway, they may not be potent inducers of the Nrf2-ARE pathway compared to Trolox.

### 3.3 Antiviral compounds in American Elderberry

Thirty-two putatively identified compounds of American elderberry juices were evaluated for their ability to inhibit HIV. Two HIV strains were used: CXCR4-dependent NL4-3 virus and CCR5-dependent BaL virus. Six of the 32 compounds tested showed significant inhibition of HIV viruses at any one concentration (Figure 6). Luteolin showed the strongest inhibition on both strains, with more than 90 percent. The second strongest inhibitor is isorhamnetin, followed by myricetin. both showed more than 50 percent inhibition of HIV viruses at a concentration of 11.1 µM. This suggested that both luteolin and myricetin can inhibit infection of HIV strains that utilize both CXCR4 and CCR5 co-receptors for their entry.

**Figure 6.**
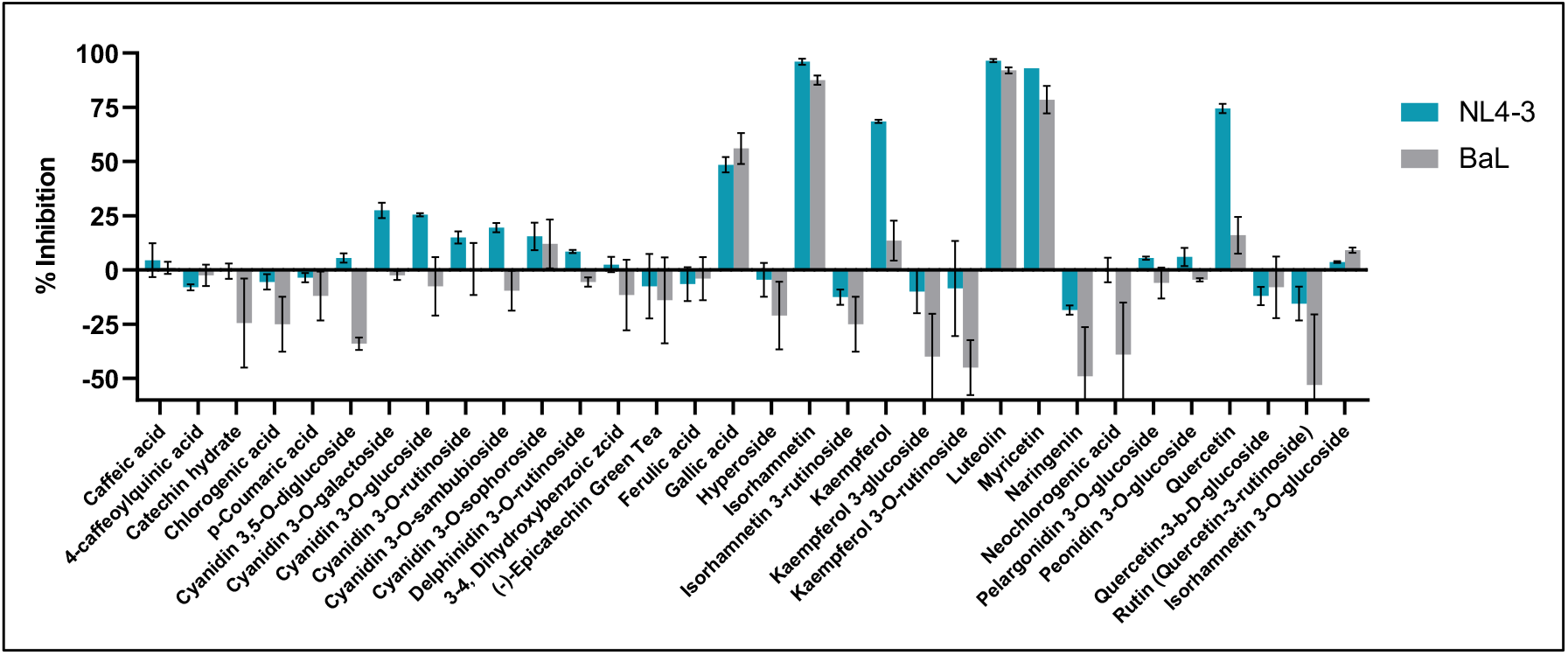
Activity of American elderberry compounds against HIV strains NL4-3 (in blue) and BaL(in grey) (n=2).

Quercetin and Kaempferol inhibited the HIV strain NL4-3 virus by more than 65 percent but didn’t significantly inhibit the HIV strain BaL virus. This suggests that quercetin and kaempferol can inhibit HIV infections that use CXCR4 as their co-receptor during viral entry. Gallic acid showed moderate inhibition of both HIV strains, with about 50 percent at a concentration of 11.1 µM. This suggested that a higher dose of gallic acid was required to inhibit the HIV strains significantly. The IC_50_ concentration of all compounds tested against HIV strains NL4-3 and BaL are presented in Supplementary Table 3.

Based on the anti-HIV assay of American elderberry putative compounds, isorhamnetin, luteolin, kaempferol, quercetin, myricetin, and gallic acid showed dose-dependent antiviral activity against CXCR4-dependent NL4-3 virus and CCR5-dependent BaL virus (Supplementary Figure 3). Isorhamnetin has the lowest IC_50_, with IC_50_ of 0.64 µM ± 0.01 against NL4-3 and 1.77 ± 0.02 µM against BaL-3. Luteolin has IC_50_s slightly higher of 1.32 µM ± 0.05 and 2.22 ± 0.69 µM against the NL4-3 and BaL viruses, respectively (Supplementary Table 3). Kaempferol has IC_50_s of 2.45 ± 0.28 µM and 8.73 ± 0.70 µM against NL4-3 and BaL viruses, respectively. Isorhamnetin, luteolin, and kaempferol were toxic to cells, with cytotoxicity concentrations of 3.042 ± 0.27 µM, 4.28 ± 0.45 µM, and 11.00 ± 0.62 µM, respectively. Quercetin showed dose-dependent antiviral activity with IC_50_ of 2.49 ± 0.24 µM against NL4-3 virus and 3.84 ± 0.56 µM against BaL virus.

Quercetin was toxic to cells at a concentration of 26.41 ± 3.91 µM. Myricetin has IC_50_ of 3.11 ±0.12 µM and 2.97 ± 1.39 µM against NL4-3 and BaL viruses, respectively. In contrast, gallic acid has IC_50_ of 6.24 ± 0.12 µM and 6.08 ± 0.04 µM against NL4-3 and BaL viruses, respectively. Gallic acid showed cytotoxicity of 80.42 ± 0.57 µM. Only myricetin showed cytotoxicity higher than 100 µm.

### 3.4 Antibacterial compounds in American elderberry

Thirty-two putatively identified American elderberry compounds were evaluated for antibacterial activity against gram-positive and gram-negative bacteria. The gram-positive bacteria strain used for testing was *Staphylococcus aureus* (Figure 7). From 32 American elderberry putative compounds evaluated for antibacterial activity against *S. aureus*, three compounds inhibit the growth of *S. aureus* by more than 50 percent at a concentration of 11.1 µM. These compounds are cyanidin 3-*O*-glucoside, cyanidin 3-*O*-rutinoside, and delphinidin 3-*O*-rutinoside. In higher doses (100 µM), five compounds showed antibacterial activity against *S. aureus*. Those compounds are cyanidin 3-*O*-sambubioside, cyanidin 3-*O*-sophoroside, pelargonidin 3-*O*-glucoside, peonidin 3-*O*-glucoside, and quercetin. Epicatechin showed more than 50 percent inhibition against *S. aureus* at a concentration 300 µM.

**Figure 7.**
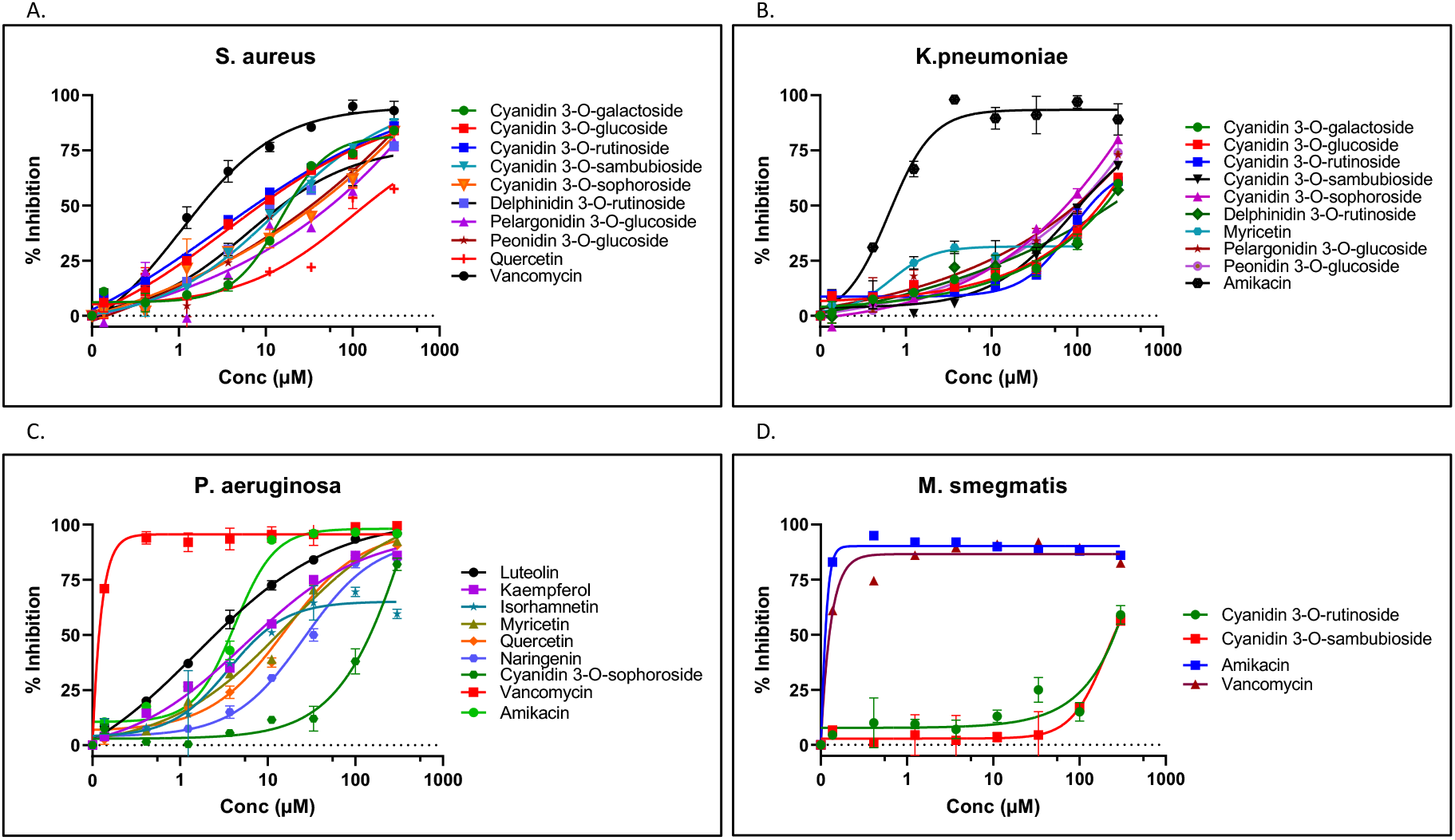
Antibacterial activity of American elderberry putative compounds against A. gram-positive bacteria *Staphylococcus aureus*, B. gram-negative bacteria *Klabsiella penumoniae*. C. gram-negative bacteria *Pseudomonas aeruginosa*, D. gram-positive and negative bacteria *M. smegmatis*. The graph showed the activity of two control compounds (Amikacin and Vancomycin) compared with two representatives of American elderberry putative compounds.

IC_50_ concentrations of six American elderberry putative compounds with significant antibacterial activity against *S. aureus* are presented in Supplementary Table 4. Six American elderberry compounds were found to have antibacterial activity against gram-positive bacteria *S. aureus*, with cyanidin 3-*O*-rutinoside having the most potent antibacterial activity (IC_50_ = 8.44 ± 0.01 µM), followed by cyanidin 3-*O*-glucoside (IC_50_ = 10.27 ± 0.52 µM). Although six compounds showed antibacterial activity against *S. aureus*, it is still less potent than Vancomycin, an antibiotic used as the positive control.

Thirty-two American elderberry putative compounds were evaluated for their antibacterial activity against gram-negative bacteria *Klebsiella pneumoniae* (Figure 7B) and *Pseudomonas aeruginosa* (Figure 7C). The majority of American elderberry putative compounds did not significantly inhibit the growth of *K. penumoniae*. Nine compounds, including cyanidin 3-*O*-sophoroside, pelargonidin 3-*O-*glucoside, and peonidin 3-*O*-glucoside, inhibit the growth of *K. penumoniae* of more than 50 percent at a concentration of 300 µM. It is less potent than the positive control antibiotic, Amikacin. The majority of American elderberry putative compounds did not significantly inhibit the growth of *Pseudomonas aeruginosa*.

Three American elderberry putative compounds, luteolin, kaempferol, and isorhamnetin, showed slightly more than 50 percent inhibition of *Pseudomonas aeruginosa* at a concentration of 11.1 µM. Five compounds showed more than 50 percent inhibition at a higher concentration (100 µM). However, they are still less potent than the positive control antibiotic Vancomycin.

Furthermore, to extend our analysis of the antibacterial activity of American elderberry putative compounds, 32 compounds were assayed against *Mycobacterium smegmatis*, bacteria with gram-positive and gram-negative characteristics. The majority of American elderberry putative compounds did not significantly inhibit the growth of *M. smegmatis* (Figure 7D). Only two compounds, cyanidin 3-*O*-rutinoside and cyanidin 3-*O*-sambubioside, showed more than 50 percent inhibition at a high concentration of 300 µM.

## 4. Discussion

American elderberry has been reported to contain health-promoting compounds^2,14,22^. In our study, we conducted an untargeted metabolomic analysis and putatively identified more than 100 metabolites (Supplementary Table 1). Based on the relative intensity, we observed the variation of these compounds in the juices across different American elderberry genotypes (Supplementary Figure 1). PLS-DA (Figure 1) and K-means clustering analysis (Figure 2) revealed that the American elderberry Ozark has a significantly different metabolomic profile than the rest of the American elderberry genotype.

To further understand the potential health benefits of these compounds, we utilize standardized HTS protocols. From over 100 putative metabolites identified, 32 metabolites with potential biological function in American elderberry juices were selected for further analysis based on their unique properties for American elderberry. This systematically comprehensive approach allowed us to capture a broad spectrum of metabolites in American elderberry juices. This approach enabled us to systematically assess the antioxidant, anti-HIV, and antibacterial activity of the 32 compounds putatively identified in American elderberry juices.

Our results showed that several putatively identified compounds in American elderberry juices exhibit strong antioxidant activities (Figure 5). Antioxidants are compounds that behave as electron scavengers to neutralize the effects of an excess of oxidants, which occurs due to an imbalance of oxidants over antioxidants that may emerge due to oxidative stress conditions^28^. An electron transfer (ET)-based assay, the CUPRAC assay, was used for our antioxidant activity study. It quantifies the ability of the compounds to counteract oxidative stress. TAC was measured by reducing chromogenic oxidant (probe), which changes color when reduced^28^. Our results indicated that 14 compounds [hyperoside (quercetin 3-galactoside), cyanidin 3-*O*-sambubioside, quercetin, cyanidin 3-*O*-galactoside, isorhamnetin, myricetin, delphinidin 3-*O*-rutinoside, cyanidin 3-*O*-sophoroside, cyanidin 3-*O*-rutinoside, cyanidin 3-*O*-glucoside, quercetin-3-b-d-glucoside (hirsutrin), cyanidin 3,5-*O*-diglucoside, neochlorogenic acid (3-caffeoylquinic acid), and gallic acid] have higher total antioxidant activity than Trolox.

Furthermore, the potential of 32 putatively identified compounds in American elderberry juices was assessed for their ability to activate the Nrf2-ARE transcriptional pathway in the HepG2 cell line. This pathway controls the gene expression of antioxidant and detoxification enzymes mediated by the ARE element in cell-based experiments^33,34^. Of all the compounds tested, four compounds (kaempferol, luteolin, isorhamnetin 3-*O-*glucoside, and quercetin) activated the Nrf2-ARE transcription pathway. None activated the Nrf2 transcription factor greater than 10-fold (Figure 5B). For a compound to be considered significant, its ability to activate the Nrf2-ARE transcription pathway must increase by more than 10-fold. Hence, despite their ability to activate the Nrf2-ARE pathway, these compounds may not be a potent inducer of the Nrf2-ARE pathway.

Our study evaluated the antiviral properties of 32 putatively identified compounds from American elderberry juices against two HIV-1 virus strains: the CXCR4-dependent NL4-3 virus and the CCR5-dependent BaL virus. Current development of anti-HIV drugs is known to target coreceptor proteins CCR5 and CXCR4, which are essential co-receptor proteins for the HIV virus to enter the host cell membrane^35^. Our findings indicated that six American elderberry putative compounds (isorhamnetin, luteolin, kaempferol, quercetin, myricetin, and gallic acid) exhibit potent antiviral activity against CXCR4-dependent NL4-3 virus and CCR5-dependent BaL virus (Figure 6) This result indicated that these compounds are potential candidates for anti-HIV inhibitors that interfere with early viral infection and replication.

To expand the therapeutic applications of American elderberry putative compounds, we evaluated the antibacterial activity against a broad spectrum of gram-positive and gram-negative bacteria. Our result suggested that nine compounds (cyanidin 3-*O*-glucoside, cyanidin 3-*O*-rutinoside, delphinidin 3-*O*-rutinoside, cyanidin 3-*O*-sambubioside, cyanidin 3-*O*-sophoroside, pelargonidin 3-*O*-glucoside, peonidin 3-*O*-glucoside, quercetin, and epicatechin) were able to inhibit gram-positive bacteria’s growth (*Staphylococcus aureus)* at one of the concentrations tested. Cyanidin 3-*O*-rutinoside has the most potent antibacterial activity (IC_50_ = 8.44 ± 0.01 µM), followed by cyanidin 3-*O*-glucoside (IC_50_ = 10.27 ± 0.52 µM). However, their ability to inhibit the growth of *S. aureus* is still less potent than Vancomycin, an antibiotic used as the positive control.

Additionally, the antibacterial activity of American elderberry putative compounds was evaluated against the gram-negative bacteria *Klebsiella pneumoniae* and *Pseudomonas aeruginosa*. The majority of American elderberry putative compounds did not significantly inhibit the growth of either of the bacteria. The 32 American elderberry putative compounds were also assayed against *Mycobacterium smegmatis*, bacteria with gram-positive and gram-negative characteristics. Most American elderberry putative compounds tested showed no antibacterial activity against *M. smegmatis*. While several American elderberry putative compounds showed promising results against gram-positive and gram-negative bacteria, their efficacy is much less potent than the common antibiotic. Further research, such as a longitudinal study involving the administration of multiple doses of the compound, is needed to optimize the antibacterial applications of these compounds.

The results from HTS assays indicated several putative compounds in American elderberry juices, including isorhamnetin, kaempferol, and luteolin, as potent bioactive compounds. Previous studies have documented a variety of pharmacological functions of those compounds. Isorhamnetin is known to have pharmacological properties such as antioxidant, anti-inflammatory, and anticancer, and it also offers cardiovascular protection^36,37^. Kaempferol is well-known for its anti-inflammatory, antibacterial, and anticancer properties^38,39^. Luteolin is a flavonoid with anticancer potential^40,41^ and has gained popularity as an antiviral agent against respiratory viruses ^42,43^. This study is the first to demonstrate the ability of Luteolin to inhibit HIV-1 virus replication.

Given the rapidly increasing popularity of American elderberry as a functional food or active ingredient in dietary supplements, our research offers valuable scientific evidence supporting its health-promoting properties. By defining the bioactive potential of American elderberry juices, we can enhance their value as functional foods and facilitate the development of innovative applications for them. As consumer interest in natural and health-promoting products grows, additional comprehensive and systematic scientific studies will be crucial for making health claims in the functional food industry.

## 5. Conclusion

This study identified 32 putative compounds from American elderberry juices. The antioxidant, antiviral, and antibacterial potentials of the putative compounds from American elderberry juices were characterized. Of 32 compounds tested, 14 were found to exhibit powerful antioxidant properties, and six showed significant antiviral activity against two strains of HIV-1 virus: CXCR4-dependent NL4-3 virus and CCR5-dependent BaL virus. Isorhamnetin, kaempferol, and luteolin are three remarkable compounds with strong antioxidant and antiviral activity. Several American elderberry putative compounds showed antibacterial activity against gram-positive bacteria *S. aureus*, with cyanidin 3-*O*-rutinoside and cyanidin 3-*O*-glucoside being the most potent. American Elderberry putative compounds didn’t show significant antibacterial activity against gram-negative bacteria *Klebsiella pneumoniae, Pseudomonas aeruginosa*, and gram-positive and gram-negative characteristics bacteria *Mycobacterium smegmatis*. These results provide scientific evidence of the health benefits of compounds identified in American elderberry juices beyond those already reported, promoting innovative application of American elderberry juices.

## 6. Acknowledgments

This work is supported by the USDA-NIFA-SCRI project 2021-51181-35860, “Moving American Elderberry into Mainstream Production and Processing”, the Missouri Department of Agriculture’s Specialty Crop Block Grant Program, and the University of Missouri’s Center for Agroforestry in cooperation with the USDA/ARS Dale Bumpers Small Farm Research Center under agreement number 58-6020-6-001. LWS is also supported by NIH NCCIH Award # 5U24AT010811-02.

## Notes

### Competing Interest Statement

The authors have declared no competing interest.

