## Supplemental Table and Fig for "Exploring American Elderberry Compounds for Antioxidant, Antiviral, and Antibacterial Properties Through High-Throughput Screening Assays Combined with Untargeted Metabolomics"

**Supplementary information**

Supplementary Table 1. The annotation of peaks isolated from American Elderberry juices

| Compounds | Formula | Adducts | Theoretical Mass | Observed Mass | Δm (ppm) | Activities | Ref |
|---|---|---|---|---|---|---|---|
| 3-4, Dihydroxybenzoic acid | C7H6O4 | $[M-H]^-$ | 154.02661 | 154.027 | 1.5 | Antioxidant, anti-inflammatory | [1] |
| 3-Furoic acid | C5H4O3 | $[M-H]^-$ | 112.01605 | 112.017 | 4.3 | Allosteric modulator | [2] |
| 3-hydroxycoumarin | C9H6O3 | $[M+H]^+$ | 162.03169 | 162.031 | -2.9 | Anticancer, anti-inflammatory, antimicrobial | [3–5] |
| 4-caffeoylquinic acid (cryptochlorogenic acid) | C16H18O9 | $[M+H]^+$ | 354.09508 | 354.095 | 1.2 | Anti-inflammatory, antioxidant, antidiabetic | [6,7] |
| 4-hydroxybenzaldehyde | C7H6O2 | $[M+H]^+$ | 122.03678 | 122.036 | -5.0 | Antioxidant, wound healing, vasculoprotective agent | [8,9] |
| 5,7-Dihydroxychromone | C9H6O4 | $[M+H]^+$ | 178.02661 | 178.026 | -2.4 | Antioxidant, antidiabetic, anticancer | [10–12] |
| 7,8-Dihydroxycoumarin (Daphnetin) | C9H6O4 | $[M+H]^+$ | 178.02661 | 178.026 | -2.4 | Neuroprotective, antioxidant, anti-inflammatory, anticancer | [13,14] |
| Allamandin | C15H16O7 | $[M+H]^+$ | 308.0896 | 308.089 | -1.3 | Antiviral, anticancer | [15–17] |
| Anethole | C10H12O | $[M+H]^+$ | 148.08882 | 148.088 | -3.6 | Antimicrobial, antioxidant, anti-inflammatory | [18,19] |
| Aurentiacin | C18H18O4 | $[M+H]^+$ | 298.12051 | 298.12 | -1.0 | Anti-inflamatory, antimicrobial | [20,21] |
| Betaine | C5H11NO2 | $[M+H]^+$ | 117.07898 | 117.078 | -4.3 | Antioxidant, anti-inflammatory | [22,23] |
| Burseran | C22H26O6 | $[M+Na]^+$ | 386.17294 | 386.173 | -0.2 | Anti-inflammatory, anticancer | [24–27] |

| | | | | | | | |
|---|---|---|---|---|---|---|---|
| Caffeic acid | C9H8O4 | [M-H]$^{-}$ | 180.04226 | 180.043 | 2.0 | Antioxidant | [28] |
| Castanospermine | C8H15NO4 | [M+H]$^{+}$ | 189.10011 | 189.1 | -2.3 | Antivirus, anticancer | [29,30] |
| Catechin | C15H14O6 | [M-H]$^{-}$ | 290.07904 | 290.079 | 0.2 | Antioxidant, anticancer, cardiovasculoprotective potency | [31,32] |
| Cearoin | C14H12O4 | [M+H]$^{+}$ | 244.07356 | 244.073 | -0.9 | Anticancer, anti-inflammatory, antimicrobial | [33–35] |
| Chlorogenic acid (5-caffeoylquinic acid) | C16H18O9 | [M+H]$^{+}$ | 354.09508 | 354.095 | 1.2 | Antimicrobial, antidiabetic, antioxidant, anti-obesity, antihypertension | [36,37] |
| Chromone | C9H6O2 | [M+H]$^{+}$ | 146.03678 | 146.036 | -3.1 | Antiallergic, anti-inflammatory, antidiabetic, anticancer, and antimicrobial | [38,39] |
| Cinnamaldehyde | C9H8O | [M+H]$^{+}$ | 132.05751 | 132.057 | -3.8 | Antimicrobial, antidiabetic, antioxidant | [40,41] |
| Citric acid | C6H8O7 | [M-H]$^{-}$ | 192.027 | 192.027 | 0.6 | Antioxidant, anti-inflammation | [42,43] |
| Cotinine | C10H12N2O | [M+H]$^{+}$ | 176.09496 | 176.094 | -4.9 | Neuroprotective | [44] |
| Coumarin | C9H6O2 | [M+H]$^{+}$ | 146.03678 | 146.036 | -3.1 | Anticancer, anti-inflammatory, antimicrobial | [3,4] |
| Cuminaldehyde | C10H12O | [M+H]$^{+}$ | 148.08882 | 148.088 | -3.6 | Antimicrobial, antioxidant, anticancer | [45,46] |
| Cyanidin 3-O-galactoside | C21H21O11 | [M+]$^{+}$ | 449.10839 | 449.108 | 1.4 | Antioxidant, anti-inflammatory, anticancer | [47] |
| Cyanidin 3-O-glucoside | C21H21O11 | [M+H]$^{+}$ | 449.10839 | 449.109 | 2.1 | Antioxidant, anti-inflammatory | [48–50] |

| | | | | | | | |
|---|---|---|---|---|---|---|---|
| Cyanidin 3-O-rutinoside | C27H31O15 | $[M+]^+$ | 595.1663 | 595.166 | 0.0 | Antioxidant, anti-inflammatory | [48,50] |
| Cyanidin 3-O-sambubioside | C26H29O15 | $[M+]^+$ | 581.15065 | 581.15 | 1.9 | Anticancer, antihypertension, antioxidant, antimicrobial | [51,52] |
| Cyanidin 3-O-sophoroside | C27H31O16 | $[M+]^+$ | 611.16121 | 611.161 | 0.2 | Anticancer, antioxidant | [53,54] |
| Cyanidin 3,5-O-diglucoside | C27H31O16 | $[M+]^+$ | 611.16121 | 611.161 | 0.2 | Antioxidant, anti-inflammatory, skin hydration effect | [55,56] |
| Dehydrozingerone | C11H12O3 | $[M+H]^+$ | 192.07864 | 192.078 | -3.7 | Antioxidant, anti-inflammatory, antidepressant, antimicrobial, antidiabetic, anticancer | [57–60] |
| Delphinidin 3-O-rutinoside (tulipanin) | C27H31O16 | $[M+H]^+$ | 611.16121 | 611.16 | 1.2 | Antioxidant, anticancer | [61,62] |
| Dimethadione | C5H7NO3 | $[M+NH4]^+$ | 129.04259 | 129.042 | -2.4 | Antiepileptic | [63] |
| Epiafzelechin trimethyl ether | C18H20O5 | $[M+H]^+$ | 316.13107 | 316.131 | -0.8 | Anticancer | [64] |
| Epicatechin | C15H14O6 | $[M-H]^-$ | 290.07904 | 290.079 | 1.3 | Antioxidant, anti-inflammatory, neuroprotective agent | [65–67] |
| Estragole | C10H12O | $[M+H]^+$ | 148.08882 | 148.088 | -3.6 | Antioxidant, anticancer | [68] |
| Ethyl p-coumarate | C11H12O3 | $[M+H]^+$ | 192.07864 | 192.078 | -3.7 | Antifungal, antioxidant | [69,70] |
| Ferulic acid | C10H10O4 | $[M+H]^+$ | 194.05791 | 194.057 | 4.3 | Antioxidant, anti-inflammatory, anticancer, antimicrobial | [71,72] |
| Gallic acid | C7H6O5 | $[M+NH4]^+$ | 170.02152 | 170.021 | 2.1 | Antioxidant, anti-inflammatory, anticancer, antimicrobial, antiviral | [73] |

| | | | | | | | |
|---|---|---|---|---|---|---|---|
| Gastrodin | C13H18O7 | $[M+Na]^+$ | 286.10525 | 286.105 | -0.6 | Neuroprotective, antidiabetic, anti-inflammatory | [74,75] |
| Herniarin | C10H8O3 | $[M+H]^+$ | 176.04734 | 176.047 | -2.7 | Anticancer, anti-inflammatory, neuroprotective | [76,76] |
| Hymechrome | C10H8O3 | $[M+H]^+$ | 176.04734 | 176.047 | -2.7 | Anticancer | [77] |
| Hyperoside (Quercetin 3-galactoside) | C21H20O12 | $[M-H]^-$ | 464.09548 | 464.096 | 0.5 | Anticancer, anti-inflammatory, antibacterial, antiviral, antidepressant | [78] |
| Indoleacrylic acid | C11H9NO2 | $[M+H]^+$ | 187.06333 | 187.063 | -2.4 | Anti-inflammatory | [79] |
| Isohopeaphenol | C56H42O12 | $[M+K]^+$ | 906.26763 | 905.265 | -1106.6 | Antioxidant, antimicrobial, antiviral | [80–82] |
| Isorhamnetin | C16H12O7 | $[M+H]^+$ | 316.0583 | 316.058 | 0.9 | Anticancer, antioxidant | [83,84] |
| Isorhamnetin 3-rutinoside (Narcissin) | C28H32O16 | $[M-H]^-$ | 624.16903 | 624.169 | 0.2 | Antiviral, antioxidant, antimicrobial, anticancer | [85–87] |
| Isorhamnetin-3-glucoside | C22H22O12 | $[M-H]^-$ | 478.11113 | 478.111 | 0.5 | Anti-inflammatory | [88] |
| Kaempferol | C15H10O6 | $[M-H]^-$ | 286.04774 | 286.048 | 1.2 | Antimicrobial, antioxidant | [89,90] |
| Kaempferol 3-glucoside (astragalin) | C21H20O11 | $[M-H]^-$ | 448.10056 | 448.101 | 1.5 | Anti-inflammatory, antioxidant, antidiabetic, anticancer, cardioprotective agent, cosmetic use | [91] |
| Kaempferol 3-O-rutinoside | C27H30O15 | $[M+H]^+$ | 594.15847 | 594.158 | 0.2 | Antioxidant | [92,93] |
| Kinetin riboside | C15H17N5O5 | $[M+NH4]^+$ | 347.12296 | 347.122 | -3.4 | Anticancer | [94,95] |
| Kuwanon L | C35H30O11 | $[M+H]^+$ | 626.17881 | 626.175 | -6.0 | Antiviral, anti-inflammatory, antimicrobial | [96–98] |

| | | | | | | | |
|---|---|---|---|---|---|---|---|
| L-tyrosine | C9H11NO3 | [M+H]$^{+}$ | 181.07389 | 181.074 | -1.9 | Antidepressant, antioxidant | [99,100] |
| L-Valine | C5H11NO2 | [M+H]$^{+}$ | 117.07898 | 117.078 | -4.3 | Amino acid | [101] |
| Lucenin 2 | C27H30O16 | [M+H]$^{+}$ | 610.15338 | 610.154 | 0.3 | Anti-inflammatory, antioxidant | [102] |
| Luteolin | C15H10O6 | [M-H]$^{-}$ | 286.04774 | 286.048 | 1.2 | Anticancer, antiviral | [103–106] |
| Monocerin | C16H20O6 | [M+H]$^{+}$ | 308.12599 | 308.125 | -3.3 | Antimicrobial | [107] |
| Moroctic acid (stearidonic acid) | C18H28O2 | [M+NH4]$^{+}$ | 276.20892 | 276.209 | -0.5 | Anti-inflammatory, anticancer, anti-acne | [108–110] |
| Myricetin (Myricitin) | C15H10O8 | [M-H]$^{-}$ | 318.03757 | 318.038 | 2.0 | Anticancer, anti-inflammatory, antidiabetic, cardio-cerebrovascular protection agent, anti-neurodegenerative | [111] |
| Myristicin | C11H12O3 | [M+H]$^{+}$ | 192.07864 | 192.078 | -3.7 | Anticancer, antimicrobial, anti-inflammatory, antioxidant | [112–114] |
| Naringenin | C15H12O5 | [M-H]$^{-}$ | 272.06847 | 272.069 | 2.3 | Anticancer, antioxidant, anti-inflammatory, antidiabetic, neuroprotective agent | [115,116] |
| Neochlorogenic acid (3-caffeoylquinic acid) | C16H18O9 | [M+H]$^{+}$ | 354.09508 | 354.095 | 0.5 | Antioxidant, antifungal, anti-inflammatory and anticarcinogenic effects | [117,118] |
| Niacin (Vit B3) | C6H5NO2 | [M+H]$^{+}$ | 123.03203 | 123.032 | -3.9 | Anti-dyslipidemic, anti-inflammatory | [119,120] |
| Niacinamide | C6H6N2O | [M+H]$^{+}$ | 122.04801 | 122.048 | -3.2 | Anti-acne, antiaging and skin improvement effect | [121,122] |
| Nutlin-3 | C30H30Cl2N4O4 | [M+K]$^{+}$ | 580.16441 | 579.165 | -1723.1 | Anticancer | [123,124] |

| | | | | | | | |
|---|---|---|---|---|---|---|---|
| p-Coumaric acid | C9H8O3 | [M-H] - | 164.04734 | 164.048 | 3.0 | Antimicrobial, antioxidant, anticancer | 125–127 |
| p-Cymene | C10H14 | [M+H] + | 134.10955 | 134.109 | -2.7 | Antioxidant, anti-inflammatory, anticancer, antimicrobial | 128–131 |
| Palmitoyl glycine | C18H35NO3 | [M+Na] + | 313.26169 | 313.262 | -0.5 | Anti-inflammatory | 132 |
| Pelargonidin 3-O-glucoside | C21H21O10 | [M+]+ | 433.11347 | 433.113 | 0.7 | Anti-inflammatory, antioxidant | 133–135 |
| Peonidin 3-O-glucoside | C22H23ClO11 | [M+H] + | 498.09289 | 498.094 | 1.6 | Anti-inflammatory, antioxidant, anticancer | 136,137 |
| Picolinic acid | C6H5NO2 | [M+H] + | 123.03203 | 123.032 | -3.9 | Anticancer, antiviral | 138–140 |
| Pyroglutamic acid | C5H7NO3 | [M+H] + | 129.04259 | 129.042 | -3.1 | Antimicrobial, anti-inflammatory, neurogenic activities | 141,142 |
| Quercetin | C15H10O7 | [M-H] - | 302.04265 | 302.043 | 0.9 | Antioxidant, anti-inflammatory, anticancer | 43,143–145 |
| Quercetin-3-b-D-glucoside (isoquercetin) | C21H20O12 | [M-H] - | 464.09548 | 464.096 | 0.9 | Antioxidant, anti-inflammatory, neuroprotective | 146–148 |
| r-Viniferin | C56H42O12 | [M+K] + | 906.26763 | 905.265 | -1106.6 | Antioxidant, anti-inflammatory, antimicrobial, anticancer | 149–151 |
| Rutin | C27H30O17 | [M-H] - | 610.15338 | 610.153 | 0.1 | Anticancer, antioxidant, anti-inflammatory | 43,146,152,153 |
| Salicin | C13H18O7 | [M+Na] + | 286.10525 | 286.105 | -0.6 | Anticancer, antioxidant | 154–156 |
| Salicylaldehyde | C7H6O2 | [M+H] + | 122.03678 | 122.036 | -5.0 | Antimicrobial | 157 |
| Salidroside | C14H20O7 | [M+Na] + | 300.1209 | 300.12 | -1.5 | Anticancer, anti-inflammation, antioxidant, antivirus, antiaging, neuroprotective | 158–160 |

| | | | | | | | |
|---|---|---|---|---|---|---|---|
| Spermine | $C_{10}H_{26}N_4$ | $[M+H]^+$ | 202.21575 | 202.215 | -2.6 | Antioxidant, anticancer, antiaging | 29,161,162 |
| Tamarixetin | $C_{16}H_{12}O_7$ | $[M+H]^+$ | 316.0583 | 316.058 | -0.9 | Anticancer, anti-inflammatory, anti-oxidation, organ protection, prevention of obesity | 163–165 |
| Tropolone | $C_7H_6O_2$ | $[M+H]^+$ | 122.03678 | 122.036 | -5.0 | Anticancer, antimicrobial | 166,167 |
| Vanilloloside | $C_{14}H_{20}O_8$ | $[M-H]^-$ | 316.11582 | 316.116 | 0.3 | Antimicrobial, antioxidant, anticancer | 168–170 |
| Vitisin C | $C_{56}H_{42}O_{12}$ | $[M+K]^+$ | 906.26763 | 905.265 | -1106.6 | Neuroprotective, antioxidant | 148,171 |

Procedures employed for the identification or putative identification[172] are comparisons with MS spectral data or accurate mass in the METLIN library.

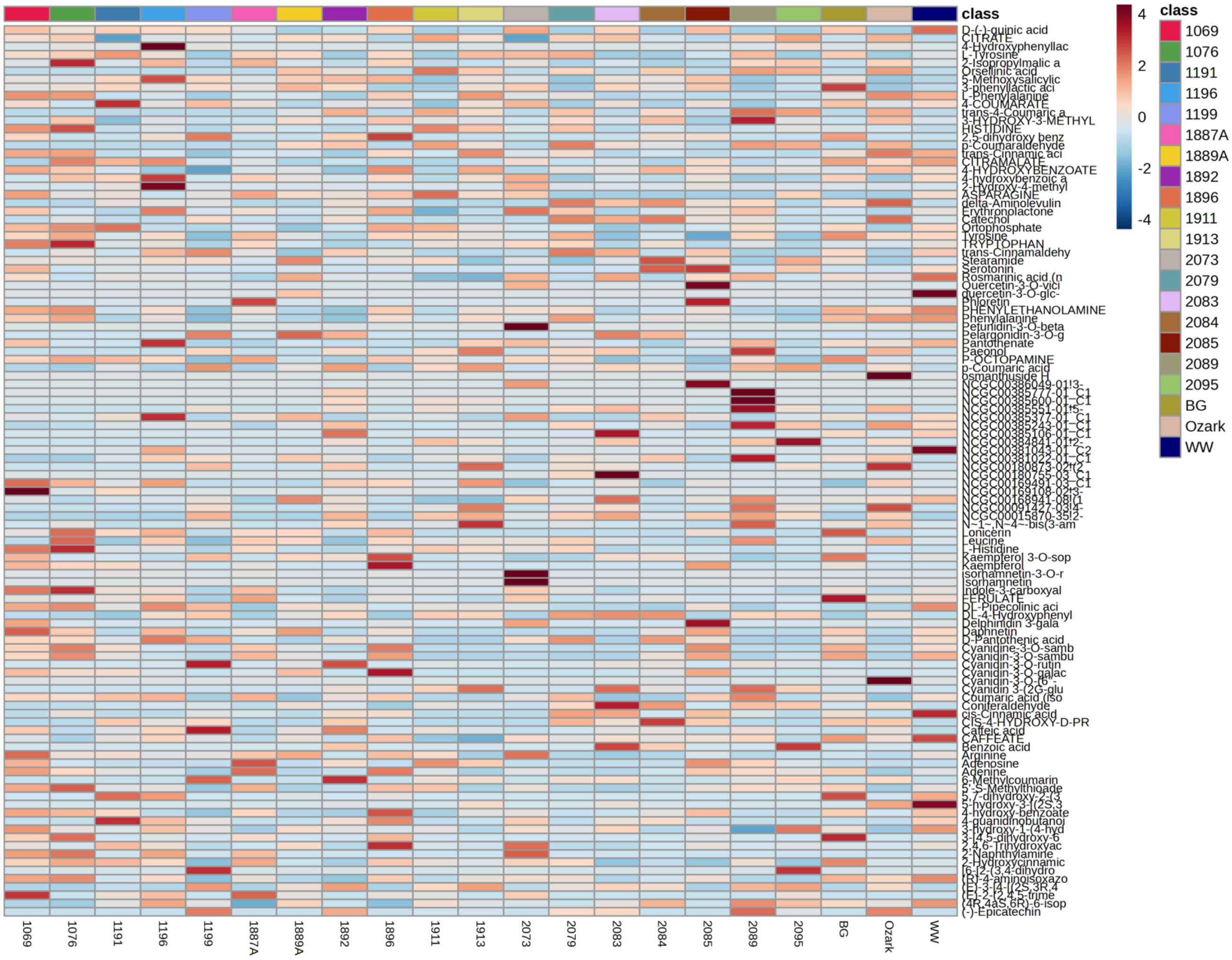


Supplementary Figure 1. The heatmap relative intensity of 108 putatively identified compounds in the juices of 21 American elderberry genotypes.

Supplementary Table 2. K-means clustering result.

| Cluster | Members |
|---|---|
| Cluster 1 | 1191 , 1191.1 , 1191.2 , 1196 , 1196.1 , 1196.2 , 1199 , 1199.1 , 1199.2 , 1889A , 1889A.1 , 1889A.2 , 1892 , 1892.1 , 1892.2 , 1911 , 1911.1 , 1911.2 , 1913 , 1913.1 , 1913.2 , 2073 , 2073.1 , 2073.2 , 2079 , 2079.1 , 2079.2 , 2083 , 2083.1 , 2083.2 , 2084 , 2084.1 , 2084.2 , 2085.1 , 2089 , 2089.1 , 2089.2 , 2095 , 2095.1 , 2095.2 |
| Cluster 2 | Ozark , Ozark.1 , Ozark.2 |
| Cluster 3 | 1069 , 1069.1 , 1069.2 , 1076 , 1076.1 , 1076.2 , 1887A , 1887A.1 , 1887A.2 , 1896 , 1896.1 , 1896.2 , 2085 , 2085.2 , BG , BG.1 , BG.2 , WW , WW.1 , WW.2 |

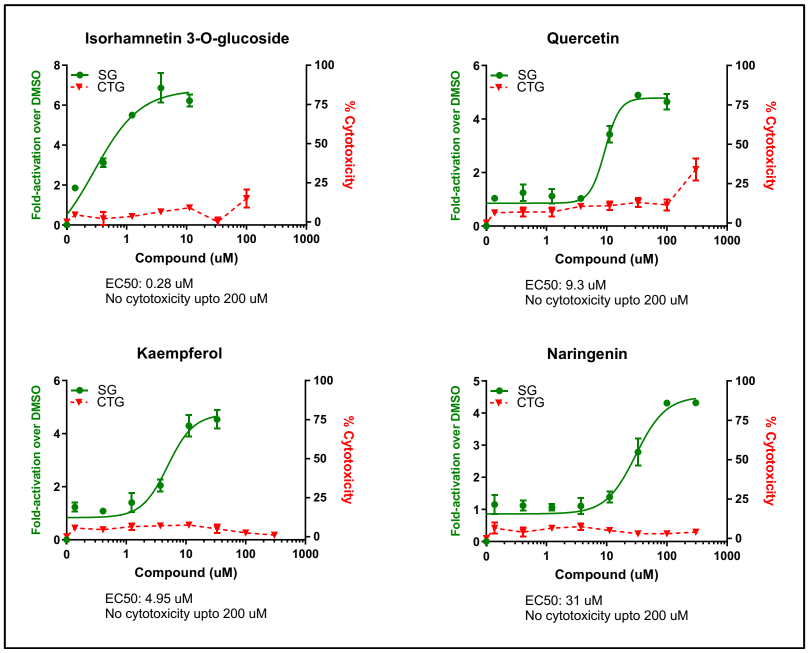


Supplementary Figure 2. Representative American elderberry compound cytotoxicity in ARE assay in HepG2 cell line (n=2).

Supplementary Table 3. $IC_{50}$ concentration of AE putative compounds against HIV strains NL4-2 and BaL

| No. | Compound name | NL4-3 | | BaL | |
|---|---|---|---|---|---|
| | | **Mean** | **SD** | **Mean** | **SD** |
| 1 | Caffeic acid | 97.14 | 13.97 | >100 | 0.00 |
| 2 | 4-caffeoylquinic acid (cryptochlorogenic acid) | >100 | 0.00 | >100 | 0.00 |
| 3 | Catechin hydrate | NI | NI | NI | NI |
| 4 | Chlorogenic acid (5-caffeoylquinic acid) | NI | NI | NI | NI |
| 5 | p-Coumaric acid | NI | NI | NI | NI |
| 6 | Cyanidin 3,5-*O*-diglucoside | NI | NI | NI | NI |
| 7 | Cyanidin 3-*O*-galactoside | 55.95 | 3.57 | >100 | 0.00 |
| 8 | Cyanidin 3-*O*-glucoside | 65.77 | 11.68 | >100 | 0.00 |
| 9 | Cyanidin 3-*O*-rutinoside | >100 | 0.00 | >100 | 0.00 |
| 10 | Cyanidin 3-*O*-sambubioside | 51.28 | 2.10 | >100 | 0.00 |
| 11 | Cyanidin 3-*O*-sophoroside | 53.31 | 5.10 | 87.37 | 2.88 |
| 12 | Delphinidin 3-*O*-rutinoside | 34.92 | 1.25 | 97.52 | 9.13 |
| 13 | 3-4, Dihydroxybenzoic acid | NI | NI | NI | NI |
| 14 | (-)-Epicatechin | NI | NI | NI | NI |
| 15 | Ferulic acid | NI | NI | NI | NI |
| 16 | Gallic acid | 6.24 | 0.12 | 6.24 | 0.12 |
| 17 | Hyperoside (Quercetin 3-galactoside) | NI | NI | NI | NI |
| 18 | Isorhamnetin | 0.64 | 0.01 | 2.22 | 0.69 |
| 19 | Isorhamnetin 3-rutinoside | NI | NI | NI | NI |
| 20 | Kaempferol | 2.45 | 0.28 | 8.73 | 0.70 |
| 21 | Kaempferol 3-glucoside | NI | NI | NI | NI |
| 22 | Kaempferol 3-*O*-rutinoside | NI | NI | NI | NI |
| 23 | Luteolin | 1.32 | 0.05 | 1.77 | 0.02 |
| 24 | Myricetin | 3.11 | 0.12 | 2.97 | 1.39 |
| 25 | Naringenin | NI | NI | NI | NI |
| 26 | Neochlorogenic acid (3-caffeoylquinic acid) | NI | NI | NI | NI |
| 27 | Pelargonidin 3-*O*-glucoside | NI | NI | NI | NI |
| 28 | Peonidin 3-*O*-glucoside | NI | NI | NI | NI |
| 29 | Quercetin | 2.49 | 0.24 | 3.84 | 0.56 |
| 30 | Quercetin-3-β-D-glucoside (hirsutrin) | >100 | 0.00 | >100 | 0.00 |
| 31 | Rutin (Quercetin-3-rutinoside) | NI | NI | NI | NI |
| 32 | Isorhamnetin 3-*O*-glucoside | NI | NI | NI | NI |

*NI = No inhibition.
**Compounds with an $IC_{50}$ concentration greater than 100μM were found to have no inhibitory on the growth of *S. aureus* (n=2).

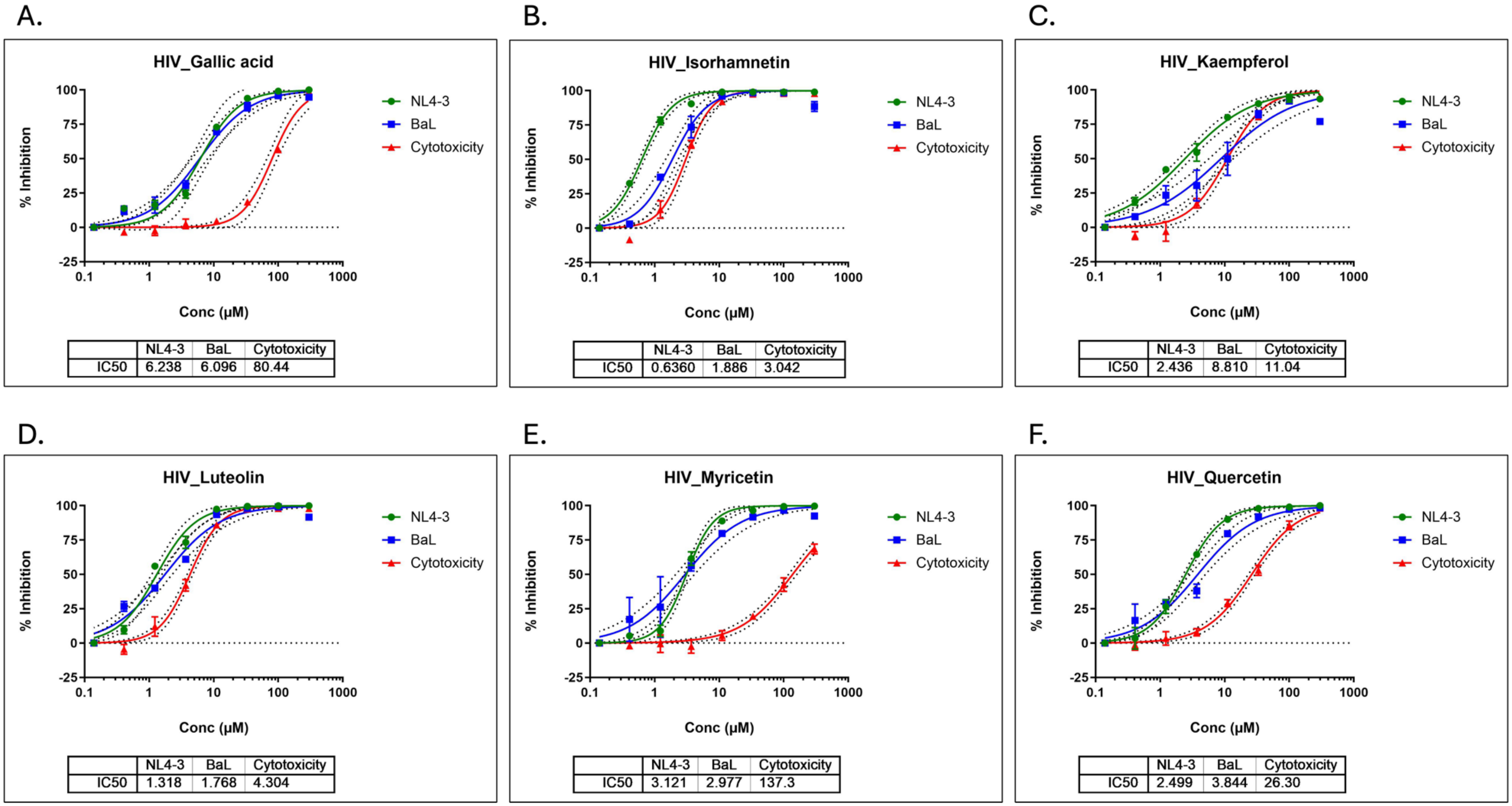

Supplementary Figure 3. Antiviral activity of American elderberry compounds against HIV strains NL4-3 and BaL (n=2).

Supplementary Table 4. The $IC_{50}$ concentration of American elderberry putative compounds against *S. aureus*

| No. | Compounds | $IC_{50}$ (μM) | SD |
|---|---|---|---|
| 1 | Caffeic acid | NI | NI |
| 2 | 4-caffeoylquinic acid (cryptochlorogenic acid) | NI | NI |
| 3 | Catechin hydrate | NI | NI |
| 4 | Chlorogenic acid (5-caffeoylquinic acid) | NI | NI |
| 5 | p-Coumaric acid | NI | NI |
| 6 | Cyanidin 3,5-*O*-diglucoside | NI | NI |
| 7 | Cyanidin 3-*O*-galactoside | 21.95 | 0.37 |
| 8 | Cyanidin 3-*O*-glucoside | 10.27 | 0.52 |
| 9 | Cyanidin 3-*O*-rutinoside | 8.44 | 0.01 |
| 10 | Cyanidin 3-*O*-sambubioside | 16.78 | 0.92 |
| 11 | Cyanidin 3-*O*-sophoroside | 32.36 | 9.19 |
| 12 | Delphinidin 3-*O*-rutinoside | 22.00 | 2.66 |
| 13 | 3-4, Dihydroxybenzoic zcid | NI | NI |
| 14 | (-)-Epicatechin | NI | NI |
| 15 | Ferulic acid | NI | NI |
| 16 | Gallic acid | NI | NI |
| 17 | Hyperoside (Quercetin 3-galactoside) | NI | NI |
| 18 | Isorhamnetin | NI | NI |
| 19 | Isorhamnetin 3-rutinoside | NI | NI |
| 20 | Kaempferol | NI | NI |
| 21 | Kaempferol 3-glucoside | NI | NI |
| 22 | Kaempferol 3-*O*-rutinoside | NI | NI |
| 23 | Luteolin | NI | NI |
| 24 | Myricetin | NI | NI |
| 25 | Naringenin | NI | NI |
| 26 | Neochlorogenic acid (3-caffeoylquinic acid) | NI | NI |
| 27 | Pelargonidin 3-*O*-glucoside | 46.45 | 5.78 |
| 28 | Peonidin 3-*O*-glucoside | 28.65 | 2.25 |
| 29 | Quercetin | >100 | 0.00 |
| 30 | Quercetin-3-β-D-glucoside (hirsutrin) | NI | NI |
| 31 | Rutin (Quercetin-3-rutinoside) | NI | NI |
| 32 | Isorhamnetin 3-*O*-glucoside | NI | NI |
| 33 | Vancomycin (Control +) | 2.16 | 0.42 |

*NI = No inhibition.

**Compounds with an $IC_{50}$ concentration greater than 100μM were found to have no inhibitory on the growth of *S. aureus* (n=2).
